# Beyond Alzheimer’s story: how an engineered molecule gained an endogenous essence

**DOI:** 10.1101/2025.09.22.677901

**Authors:** Arthur T. Kopylov, Ruslan Yu. Yakovlev, Aleksei V. Trukhin, Yuri M. Poluektov, Anastasia A. Anashkina, Sergey A. Kozin

**Author notes:** Correspondence (S.A.K.).

## Abstract

A hallmark of Alzheimer’s disease (AD) is extra-neuronal protein aggregates formed mainly by the amyloid-β peptide (Aβ), that are deposited inside the brain as amyloid plaques. The engineered peptide Ac-His-Ala-Glu-Glu-NH_2_ (HAEE) suppresses the formation of amyloid plaques *in vivo* and is being tested as an anti-amyloid drug candidate. Here, by using a quantitative mass-spectrometry method we discover that HAEE peptide represents a normal endogenous component of human blood plasma. Moreover, the HAEE level has been significantly reduced in patients with a clinical diagnosis of AD (n=200) as compared with the control participants with no clinical diagnosis of cognitive impairment (n=150). Thus, endogenous HAEE may be considered as a potential blood-based biomarker in the diagnosis of Alzheimer’s disease, while exogenous HAEE may serve as a drug in AD-modifying therapy.

## INTRODUCTION

Alzheimer’s disease (AD) is the most common form of primary degenerative dementia of late age (Livingston et al., 2020), which is clinically characterized by a gradual, barely noticeable onset in presenile or old age, a steady progression of memory disorders and higher cortical functions up to the total collapse of intelligence and mental activity (Long and Holtzman, 2019).

Currently, AD is defined by the following neuropathological profile: (1) deposition of amyloid-β peptide (Aβ) aggregates in the form of diffuse and amyloid (neuritic, senile) plaques and (2) the presence of intraneuronal neurofibrillary tangles and neuropil filaments (in dystrophic neurites) comprised of aggregated hyperphosphorylated tau protein (Jagust et al., 2019).

Depending on the stage of the disease, the following idealized therapeutic intervention regimen for the treatment of AD has been proposed: (1) inhibition of Aβ formation and/or aggregation; (2) removal of Aβ aggregates and/or neutralization of the toxicity of such aggregates; (3) blocking the extracellular spread of tau protein; (4) protection from the toxic effects of tau protein; (5) compensation for cellular dysfunctions; (6) regeneration of affected neuronal cells; (7) symptomatic therapy (Golde et al., 2018).

Translational studies—including experimental animal and human neuropathological, genetic, and *in vivo* biomarker-based evidence—support a descriptive hypothetical model of AD pathophysiology characterized by the upstream brain accumulation of Aβ species and plaques, which precedes spreading of tau, neuronal loss and ultimately clinical manifestations by up to 20–30 years (Hampel et al., 2021). Thus, Aβ aggregation may be a key event in initiating the AD pathological cascade, while neuroinflammation and tau protein accumulation may be the main drivers of neurodegeneration (Long and Holtzman, 2019).

Very recently three anti-Aβ antibodies - aducanumab, lecanemab, and donanemab (Budd Haeberlein et al., 2022; Mintun et al., 2021; van Dyck et al., 2023) - have been approved by the US Food and Drug Administration for treatment initiation in early AD comprising patients with mild cognitive impairment and mild dementia due to AD with Aβ -based diagnostic confirmation using amyloid positron emission tomography or cerebrospinal fluid biomarkers. These new disease□modifying treatments act by lowering brain amyloid and demonstrate the clinical benefits of anti-amyloid therapy (Hardy and Mummery, 2023). It is important to note that current anti-amyloid therapies that bind to and clear amyloid plaques have also prompted safety concerns (Kepp et al., 2023).

Several experimental observations suggest that Aβ might interact with zinc ions during AD pathogenesis: (i) amyloid plaques contain excessively high zinc ion level (Lovell et al., 1998) (ii) zinc binds to Aβ and elicits its rapid aggregation (Bush et al., 1994), presumably due to modulating Aβ conformational transformation and populational shift in the equilibrium of Aβ polymorphic states (Miller et al., 2010); (iii) post-mortem brain samples from the patients with AD were found to have zinc-rich areas overlapping with the sites of amyloid plaque formation (Miller et al., 2006); (iv) the AD-affected brain areas are enriched with zinc-containing axons, while the less affected brain regions contain only low levels of zinc-containing axons (Suh et al., 2000); (v) studies of the impact of the genetic ablation of ZnT3 in the Tg2576 mouse model of Alzheimer’s disease have provided evidence that synaptically released zinc underlies amyloid pathology (Lee et al., 2002).

Based on the above, we hypothesized that abnormal interactions with zinc ions are an important prerequisite for Aβ aggregation and amyloid plaque formation in the pathogenesis of Alzheimer’s disease, and therefore a logical step was to create a molecular agent that could inhibit Aβ interactions with zinc ions and thus be a potential anti-amyloid drug.

Studies on the molecular mechanism of interactions between zinc ions and Aβ (reviewed in (Kozin, 2023)) have revealed that the central role in these interactions is played by the 11-EVHH-14 region, which: (1) is the primary zinc-binding center in Aβ monomers; (2) forms zinc-dependent interfaces in Aβ dimers and/or oligomers; (3) keeps its backbone conformation practically unchanged in both zinc-free and zinc-bound states (PDB IDs: 1ZE7, 1ZE9, 2MGT). Such extraordinary structure and function features make the EVHH site of Aβ an excellent drug target.

Molecular docking and molecular dynamics allowed us to establish that the tetrapeptide Ac-His-Ala-Glu-Glu-NH_2_ (HAEE), due to its steric and charge complementarity, is the best candidate of protein nature for the formation of a stable complex with the EVHH site of Aβ both in the absence and in the presence of a zinc ion at the intermolecular interface (reviewed in (Kozin, 2023)). Thus, engineered peptide HAEE (PubChem CID: 56971578) has the potential to both prevent the oligomerization of free Aβ molecules in biological fluids and destroy insoluble Aβ aggregates that form during the pathogenesis of Alzheimer’s disease.

In preclinical studies of the synthetic tetrapeptide HAEE as an anti-amyloid drug candidate for the treatment of Alzheimer’s disease, the following results have been obtained. Serial intravenous injections of HAEE into AβPP/PS1 double transgenic mice result in a significant reduction in amyloid load in the brains of experimental animals (Tsvetkov et al., 2015). The HAEE peptide, when administered repeatedly to rats for 28 days, did not cause visible toxic effects and abnormalities in animal behavior in doses of 300 μg/kg/day (Zolotarev et al., 2021). The pharmacokinetic parameters of HAEE were established using uniformly tritium-labeled HAEE, and pharmacokinetic data provided evidence that HAEE crossed the blood–brain barrier (Zolotarev et al., 2021). Based on molecular modeling, the role of LRP1 in receptor-mediated transcytosis of HAEE was proposed (Zolotarev et al., 2021). Additionally, in transgenic nematodes there was found the critical role of HAEE in blocking Zn-dependent aggregation of endogenous Aβ molecules in the presence of Aβ isomerized at Asp7 residue (Mitkevich et al., 2023). Altogether, the results obtained indicate that the anti-amyloid effect of HAEE most likely occurs due to its interaction with Aβ species directly in the brain and depends on its ability to inhibit zinc-mediated interactions involved in Aβ aggregation.

Next, in preparation for the proposed clinical trials of the HAEE peptide, we developed a quantitative method for monitoring HAEE in human blood plasma probes. The concentration of HAEE in blood plasma is measured using a method that combines the separation of high-performance liquid chromatography and registration on a mass detector (HPLC-MS) of the parent quasi-molecular ion and fragment ions after preliminary extraction on an ion-exchange solid-phase carrier of the deproteinized fraction of biological material (blood plasma).

## RESULTS

### HAEE is a human blood plasma endogenous component

Quite unexpectedly, we found that all plasma probes from healthy volunteers (n=150) contained endogenous HAEE at concentrations of 360±17.9 pg/mL, p <0.00001 (Table 1, Figure 1A). It is worth noting that the HAEE level is like the total Aβ level in human plasma which is 469.12±141.09 and 563.98±178.47 pg/mL for non-demented volunteers and AD patients, respectively (Roher et al., 2009). To our best knowledge, this is the first time that an artificially created tetrapeptide has been found to be an endogenous component in the human body.

**Table 1.**
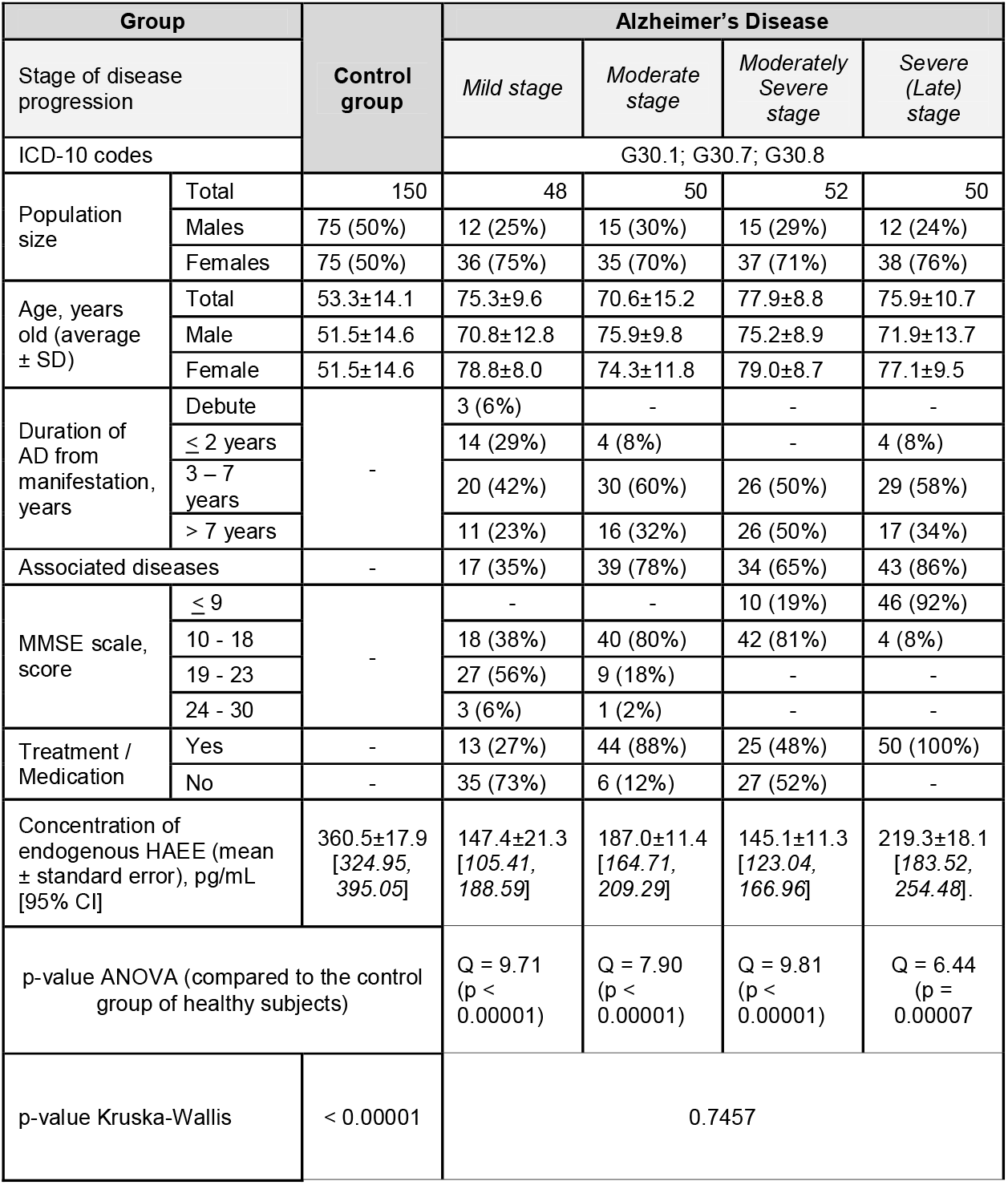
Measured concentration of the endogenous HAEE (pg/mL) and basic anthropometric and demographic data collected for patients with AD and healthy volunteers participated in the study.

**Figure 1.**
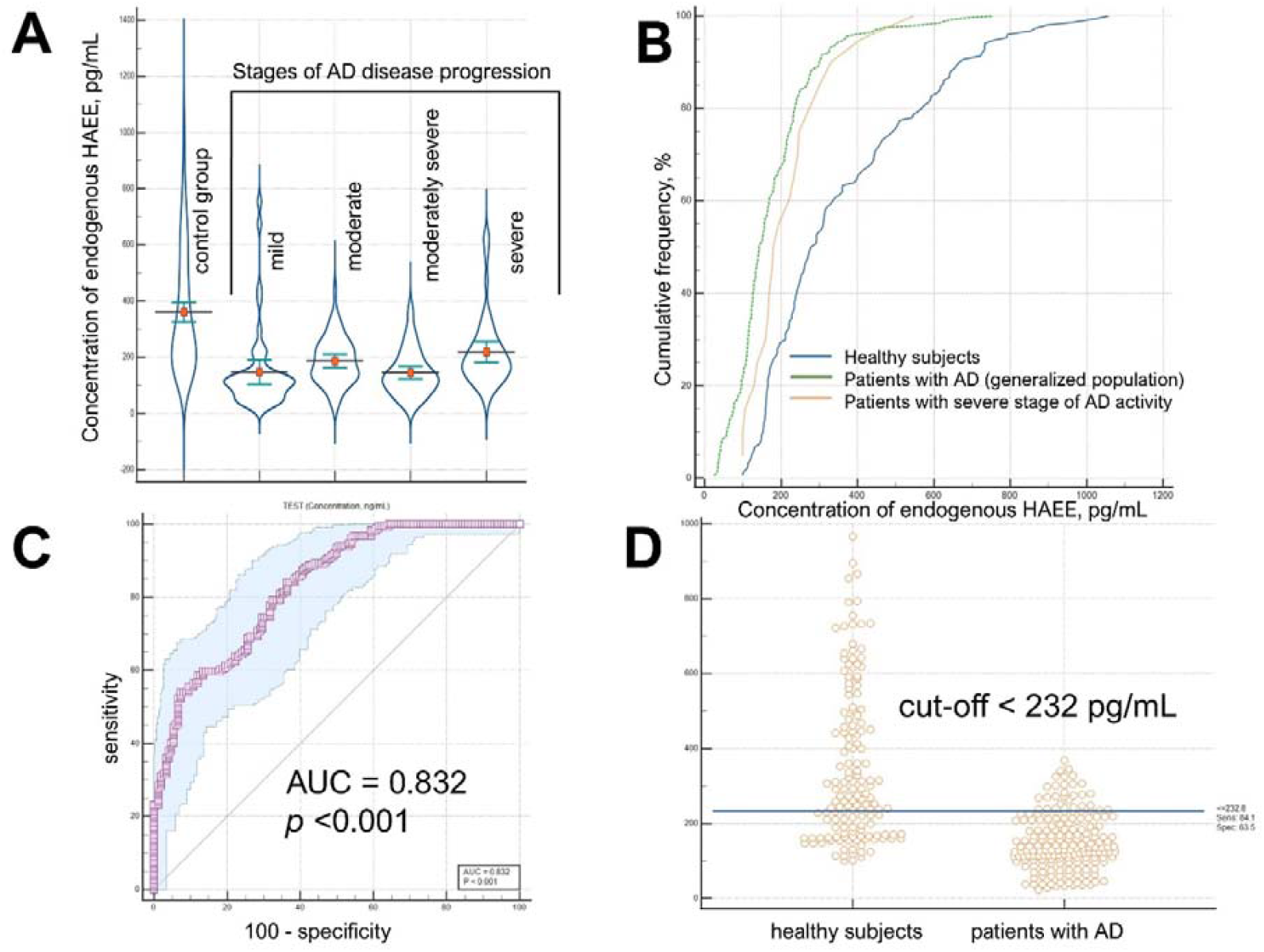
Distribution of HAEE amid patients clinically diagnosed with Alzheimer’s disease (AD), stratified by the stage of the disease activity, and a group of healthy volunteers. The median in group with mild stage was 111.9 pg/mL, the mean value was 147.4 pg/mL, the standard error of the sample was 21.3 pg /mL; in the group with moderate stage the median concentration was 176.5 pg/mL, the mean value was 187.0 pg/mL, the standard error of the sample was 11.4 pg/mL; in the group with moderate-severe stage the median concentration was 125.2 pg/ml, the mean value was 145.1 pg/mL, the standard error for the sample was 11.3 pg/mL; in the group with severe stage, the median concentration was 183.5 pg/mL, the mean value was 219.3 pg/mL, the standard error for the sample was 18.1 pg/mL. The generalized population of patients Alzheimer’s disease (all stages) showed the median concentration 143.1 pg/mL, the mean value 175.0 pg/mL, the standard error 8.2 pg/mL. The control group (healthy subjects) demonstrated the median concentration at 285.0 pg/mL, the mean value at 360.5 pg/mL, and the standard error of 17.9 pg/mL; (A). Ranged distribution of HAEE concentration in healthy subjects, patients with Alzheimer’s disease (generalized population) and patients with Alzheimer’s disease with severe stage. Up to 21% of the measured concentration values ranged up to 200 pg/mL are characterized by homoscedasticity thereafter a pronounced heteroscedastic distribution is observed for almost 79% of the entire general sample of study participants (B). ROC curve graph for a sample of 350 patients grouped by the criterion the established diagnosis of Alzheimer’s disease. The calculated AUC=0.832 with a p <0.001. The sensitivity of the test was 84.13%, the specificity of the test was 63.51%, the Youden J index was 0.4764 (C). The distribution of the measured concentration of HAEE in plasma samples of healthy subjects and patients with AD with the established cut-off level based on the ROC curve (D). Images were created using the Med program Calc version 23.0 (https://www.medcalc.org/).

A possible source of endogenous HAEE can be suggested based on the following considerations. Most physiologically active peptides including Aβ are formed from precursor proteins by the targeted action of certain proteases and/or peptidases. Many endogenous peptides undergo enzymatic acetylation at the N-terminus. In contrast, the number of endogenous peptides amidated at the C-terminus is very limited, and C-amidation occurs due to enzymatic conversion of the glycine residue at the C-terminus into an amide group (reviewed in (Kumar et al., 2016)). Since endogenous HAEE has both an N-acetyl and a C-amide group, the precursor protein must contain the pentapeptide fragment HAEEG in its amino acid sequence. The HAEEG region is found in six human proteins: (1) ciliogenesis-associated TTC17-interacting protein (*CATIP*gene); (2) reticulophagy regulator 1 (*RETREG1* gene); (3) flavin-containing monooxygenase 3 (*FMO3* gene); (4) sphingomyelin phosphodiesterase 3 (*SMPD3* gene); (5) G-patch domain-containing protein 8 (*GPATCH8* gene); (6) voltage-gated calcium channel subunit N-type alpha-1B (*CACNA1B* gene). So, at least one of these proteins may be a potential source of endogenous HAEE.

### The deficiency of endogenous HAEE is a potential blood-based biomarker for Alzheimer’s disease

The HAEE peptide has been designed as the best low-molecular-weight protein agent specifically interacting with the 11-EVHH-14 region of Aβ, and the existence of corresponding non-covalent complexes between HAEE and Aβ was demonstrated *in vitro* (Barykin et al., 2020; Mitkevich et al., 2023). Based on the fact that HAEE and Aβ are endogenous components of blood and are present there in comparable concentrations, we have hypothesized: (1) HAEE complexes with Aβ may be present in the blood; (2) these complexes may have physiological significance; (3) deficiency of complexes between HAEE and Aβ may be associated with the development of pathological conditions in Alzheimer’s disease; (4) deficiency of Aβ and HAEE complexes in Alzheimer’s disease may be caused only by HAEE deficiency, since the concentration of total Aβ in plasma is approximately the same in both healthy individuals and patients at different stages of Alzheimer’s disease (Roher et al., 2009).

The above hypothesis has been tested on blood samples obtained from patients clinically diagnosed with Alzheimer’s disease (n=200). Using the analytical method developed by us, it was found that in generalized population patients with a clinical diagnosis of AD the concentration of endogenous HAEE is 175±8.2 pg/mL, i.e. significantly reduced (almost twofold) compared to healthy volunteers; notable, there is no significant difference (p=0.746) in HAEE concentration between group of patients with various stages of disease activity (Table 1, Figure 1A and 1B). Thus, our data (Table 1) indicate that HAEE deficiency is a potential blood-based biomarker (BBM) for Alzheimer’s disease.

The receiver operating characteristic returned AUC=0.832±0.0215 (95% CI [0.787; 0.870]; Table 2, Figure 1C) for the studied population with high sensitivity (84.1%) and successes specificity (73.5%) with the recommended cut-off concentration of HAEE at 232 pg/mL (Table 2, Figure 1D) at 82.6% accuracy. The estimated positive prognostic value of HAEE has been shown to be 75.42% (95% CI [71.09; 79.29]) with a relatively high positive likelihood ratio.

**Table 2.** Estimated diagnostic value of quantitative assessment of HAEE in blood plasma for differentiating of healthy subjects from patients clinically diagnosed with Alzheimer’s disease.

| Character | Calculated Value |
| --- | --- |
| Sensitivity | 84.1 % |
| Specificity | 63.50% |
| Accuracy | 82.65% |
| Index - J (Juden) | 0.4764 |
| Recommended limit for a positive test | ≤ 232 pg/ml |
| Positive predictive value | 75.42%; 95% CI [ 71.09; 79.29 ] |
| Negative predictive value | 75.04 % ; 95% CI [ 67.93; 81.01 ] |
| Positive likelihood ratio | 2.45 |
| Negative likelihood ratio | 0.36 |
| True prevalence in the population | 56.08% |
| Area under the curve (AUC) | 0.832±0.0215; 95%CI [ 0.787; 0.870 ] |

## DISCUSSION

The discovery that a previously engineered peptide HAEE aimed at reducing amyloid burden in Alzheimer’s disease is an endogenous component of human blood plasma is noteworthy.

However, equally important is the fact that plasma levels of this component are reduced in patients with a clinical diagnosis of AD, and therefore HAEE deficiency may play a role as a potential blood-based biomarker for Alzheimer’s disease.

Current clinically available BBMs for AD can be categorized into various classes (Schöll et al., 2024). The first class includes biomarkers associated with the presence of two classic pathological hallmarks of Alzheimer’s disease: amyloid-β peptides (low Aβ42-to-Aβ40 ratio [Aβ42/40]) and elevated phosphorylated tau (p-tau) levels. The second class of blood biomarkers for Alzheimer’s disease is associated with neuronal loss, neurodegeneration, or synaptic degeneration (elevated Neurofilament light chain (Nfl) levels). The third class includes processes related to neuroinflammation mediated by glial cells (elevated Glial fibrillary acidic protein (GFAP) levels). It is obvious that HAEE deficiency as a potential AD BBM cannot be classified into any of the above-mentioned classes.

The HAEE deficiency in the blood of patients clinically diagnosed with Alzheimer’s disease compared to healthy volunteers shown in the present study (Table 1) may be caused by suppression of normal HAEE generation and/or excessive HAEE consumption due to the body’s response to various types of stress associated with the pathogenesis of Alzheimer’s disease. If we assume that HAEE is a natural partner of Aβ, then the observed HAEE deficiency in the blood of patients with AD may negatively affect the ability of circulating Aβ to serve its physiological function and/or to be in a monomeric state. So, such deficiency may be associated with the pathogenesis of Alzheimer’s disease. Moreover, one can rationally assume that the HAEE itself may play a role as constitutive neuroprotector, whose deficiency would switch on AD pathogenesis.

Overall, the results presented in this work indicate that HAEE deficiency is a potential blood-based biomarker for Alzheimer’s disease and that compensating for such deficiency, for example by administering synthetic HAEE to Alzheimer’s disease patients, may prove to be effective disease-modifying treatment.

### Limitations of the study

A major limitation is the AD cohort diagnosis, which relies solely on clinical guidelines without clear evidence of imaging or pathological biomarkers, a crucial standard for accurate diagnosis and differentiating AD from other forms of dementia. Thus, the patients under the study can represent not only Alzheimer’s Disease but also Alzheimer’s Disease Related Dementias (ADRD). While AD is the most common dementia diagnosis, ADRDs share many cognitive and pathological features with and can be difficult to distinguish from AD by using the MMSE (Mini-Mental State Examination) score.

### DATA AND MATERIAL AVAILABILITY

All data are available in the main text or the supplementary materials. The synthetic standards of HAEE (light- and [^13^C]-labeled molecules) are available upon request to A.V.T. for academic, noncommercial studies through regular material transfer agreements.

### Lead contact

Further information and requests for resources and reagents should be directed to and will be fulfilled by the lead contact, Sergey A. Kozin

## ACKNOWLEDGMENTS

We could not have written this manuscript without the support of our colleagues Sergey V. Gabestro from Lifemission, LLC (Moscow, Russia), and Eugene V. Myachin from Proteonika, LLC (Moscow, Russia). They both encouraged us to take on the project — one we would not have carried to its end were it not for their sharp, yet always generous, criticism. The research was organized and funded by Lifemission, LLC (Moscow, Russia).

## AUTHOR CONTRIBUTIONS

Conceptualization: S.A.K., A.T.K., RYY, Y.M.P., A.A.A.. Methodology: A.T.K., A.V.T.. Formal analysis: A.T.K., A.A.A., Y.M.P.. Project administration: S.A.K., A.T.K.. Funding acquisition: S.A.K.. Supervision: S.A.K.. Writing – original draft: S.A.K., A.T.K., A.V.T..

## SUPPLEMENTAL INFORMATION

**Supplementary figures** (Figs. S1 to S6)

## MATERIALS AND METHODS

### Chemicals and Reagents

Acetonitrile HPLC Gradient grade (Scharlau S.L., Spain), Formic acid 98%+ (Sigma-Aldrich, Germany); Heptafluorobutyric acid (Fisher Scientific, Hampton, NH), Methanol HPLC and spectroscopy grade (J.T. Baker, the Netherlands), Ammonia hydroxide 28-30% (Sigma-Aldrich, St. Louis, MO, USA). Analytical [^13^C_2_]-labeled (^13^C_2_C_19_H_31_N_7_O_9_ × 0.84TFA salt, purity >98%, order no. SL104559) and non-labeled (C_21_H_31_N_7_O_9_ purity >98%, order no. SL104171) standards of *Acetyl*-HAEE-*Amide* (HAEE) were synthesized by Synthon-Lab Ltd (Saint-Petersburg, Russia), and the structure was confirmed by the LC-UV-MS and ^1^H NMR analysis (Supplementary Figures S1-S6).

### Ethical Consideration

All samples were purchased from the National Bio Service (NBS, Ltd., Saint Petersburg, Russian Federation) as the main Russia operator of blood samples collection, transportation and provide a compliance of ethical consideration. All subjects from the assay group and the control group gave their written informed consent to participate in the study before the screening and samples collection in accordance with the WMA Declaration of Helsinki on Ethical Principles for Medical Research Involving Human Subject (revised Fortaleza, 2013 and revised Helsinki, 2024) and Good Clinical Practice Guidelines approved by the Russian Ministry of Health.

Subjects were enrolled to participate in the study from four clinical centers located in Moscow, Saint Petersburg, Tomsk and Nizhny Novgorod. All patients in the study group were enrolled in accordance to the current Clinical guidelines for cognitive disorders of old and senile age, Clinical guidelines developed by the Russian Association of Gerontologists and Geriatricians, and the Russian Society of Psychiatrists (revised in 2020 year).

### Population and Demography

Totally, 350 subjects participated in the observational study, of them 150 healthy volunteers with an average age of 53.3±14.1 years old were included in the control group and 200 subjects with an average age 75.6±10.3 were included in the study group of patients clinically diagnosed with Alzheimer’s disease (AD). The latter group was subdivided into four groups by the criterion of disease severity according to the recommendations of Johns Hopkins University and Alzheimer’s Association. The study group (200 subjects) comprised of 48 patients with mild stage of AD, 50 patients with moderate (middle) stage, 52 patients with the moderately sever decline and 50 patients with severe (late) stage of AD. The inclusion criteria for the subjects with AD included the presence of clinically established and monitored diagnosis of Alzheimer’s disease with the read ICD-10 codes of G30.1, G30.7, G30.8 and/or F00.0, F00.1, F00.2, and F00.9. The exclusion criteria included a history or presence of human immunodeficiency virus (HIV), hepatitis B and C, or damage of central nervous system caused by infection, trauma and/or brain injury. Subjects of the control group should also meet the criteria of absence of severe or chronic mental or neurological pathology, organic damage of the central nervous system, cardiovascular disease, absence of subjective and objective cognitive impairments at the time of inclusion in the study, concomitant decompensated somatic or neurological pathology (including epilepsy). Smoking and alcohol consumption were also recorded and recognized in associated records for each subject.

All subjects in the study group had a previous clinical history of AD (G30.1, G30.7 and G30.8 by the ICD-10) for at least two years from manifestation and the prevalence of population (87%) were monitored for 3-7 years and more. The average age at established diagnosis was 72.3±9.9 years old. Only three people from the subgroup of mild stage were enrolled as patients with normal outward behavior or preclinical stage of AD. The majority of subjects in the study group had associated diseases including but not limited to ischemic heart disease, gastritis, atherosclerosis, type 2 diabetes mellitus, etc. Most of the patients (92%) had no heredity loading of the familiar AD, whereas for the rest 8% data of heredity is unknown. The MMSE score varied from 9 and less in moderately sever and sever stages to 24 in subgroups with moderate and mild stages, nevertheless 52% of subjects scaled by MMSE within 10-18 scores. Up to 66% of the total population of patients with AD had various scheme of therapy considering associated diseases, if else, and prescribe medication including Rivastigmine, Memantine, Haloperidol, Galantamine, Amlodipine, Lisinopril, etc.

### Samples Collection

Venous blood plasma was collected in EDTA-2K vacuum tubes (Guangzhou Improve Medical Instruments Co., Ltd, Chine) between 8 a.m. and 11 a.m. hours for 2023-2024 years. Blood samples were centrifuged at 3,500 *g* for 5 minutes at room temperature in 30-40 minutes once after blood collection. The resulting plasma was split in 4-5 enumerated aliquots by 1 mL. Four to five aliquots obtained from each subject were packed into batch and designated by the subject’s clinical history ID, and frozen at -20°C for storage upon delivery. Batches of samples were occasionally delivered on dried ice for 1 – 3 days from clinical centers in Moscow, Saint Petersburg, Tomsk and Nizhny Novgorod. The custody of samples collection, pre-analytical blood preparation, tracking the changes of temperature during the storage and delivery was recorded and supported under the auspice and responsibility of the NBS service. Once samples were delivered in the laboratory, they were stored at -20°C until samples preparation. Sample preparation was accomplished using only unique aliquot from the patient’s batch to avoid the impact freeze-thaw cycle.

### Instrumentation

The analysis was carried out on a high-resolution quadrupole time-of-flight (Q-TOF) mass spectrometer (Xevo G2-XS Q-TOF, Waters, Inc., the UK) coupled with an Acquity UPLC H-Class Plus chromatography system (Waters, Inc., Singapore) and on a triple-quadrupole (QQQ) mass spectrometer (G6490A, Agilent, Inc., Singapore) enhanced by the Infinity II 1290 UPLC system (Germany). Both instruments operated in a positive ionization mode.

The Q-TOF mass spectrometry was equipped with a Z-spray™ ionization source with a capillary voltage of 3 kV and sampling cone voltage was adjusted to 21 V with the source offset of 37 V. The cone gas flow was set to 50 L/h, and the source temperature was set to 150°C whereas the desolvation gas flow was 700 L/min with the temperature of 440°C. The Step Wave was manually adjusted to achieve high sensitivity and selectivity to the target signal as following: for the Step Wave-1 wave velocity 400 m/s, wave height 9.5 V; for the Step wave-2 wave velocity 435 m/s and wave height 16.5 V; for the Step Wave DC was tuned step wave-2 offset to 18, difference aperture-1 to 4.2 V and difference aperture-2 to 0.9 V; for the source ion guide adjusted wave velocity to 367 m/s and wave height to 1.2 V.

The QQQ mass spectrometer was equipped by the Jet Stream™ ionization source with a capillary voltage of 3.2 kV and nozzle voltage of 600 V. The desolvation gas flow was set to 13 L/min with a temperature of 260°C, and the sheath gas flow was set to 8 L/min with a temperature of 320°C. The high-pressure RF voltage was adjusted to 135 V and the low-pressure RF voltage was adjusted to 60 V.

Both target molecules (intact and [^13^C_2_]-labeled HAEE) were registered and detected in a TOF-MRM (multiple reaction monitoring) mode as protonated ions with *m/z* = 526.2256 and *m/z* = 528.2329 for intact and [^13^C_2_]-labeled HAEE, respectively, by four transitions:526.2256→180.0768 (b1-ion as ion-quantifier);152.0818 (a1-ion as qualifier); 251.1139 (b2-ion as qualifier); 380.1565 (b3-ion as qualifier) and 528.2329→182.0814 (b1 ion-quantifier); 154.0893 (a1-ion); 253.0855 (b2-ion); 382.1641 (b3-ion) for the [^13^C_2_]-labeled HAEE. The scanning time on Q-TOF consisted of 418 msec and ions were surveyed within a range from 100 to 600 m/z. The selected transitions resulted after decomposition of the parent ion collision cell under argon with a collision energy ramping between 19-36 eV. The analysis was accompanied by mass correction using the lock-mass of warfarin (*m/z* = 309.1127) injected for 0.1 msec every 10 s and averaged by three scans isolated within 0.1 Da window. When the analysis was transferred to QQQ instrument, the full duty cycle consisted of 277 msec and the fragment ions were produced under the nitrogen with a collision energy 23 – 28 eV. The total number of concurrent transitions was eight and the dwell time per one transition was 33.43 msec. All the transitions passed from the collision cell to the third quadrupole with a cell accelerating voltage of 7 V.

The substances of interest were separated on an Acquity™ Premier HSS T3 column (2.1 × 150 mm; Waters, Inc., Ireland) heated to 50°C, if analyzed on the Q-TOF mass spectrometer. Molecules were separated in a gradient of mobile phase A (water) and mobile phase B (acetonitrile) both supplemented with 0.1% formic acid at a flow rate of 0.3 mL/min in the following step-wise gradient scheme: 0 to 0.5 minutes is 2-30% of B, then hold 30% of B to 1.3 minutes; then rapid increase to 78% of B at 1.9 minutes and keeping isocratic 78% of B to 2.8 minutes; then rapid increase to 92% of B at 3.2 minutes and washing the column to 4.2 minutes at 0.45 mL/min flow rate; then descend to 2% of B at 4.5 minutes and reconstitute the column in initial conditions at 0.3 mL/min for the next 2.5 minutes.

If signal recorded on QQQ instrument, molecules were separated on a Polaris™ Amide-C18 column (2.0 × 150 mm; Agilent, Inc, USA) heated at 35°C at flow rate of 0.3 mL/min in a gradient of mobile phase A (water) and mobile phase B (methanol) both supplied with 0.08% of formic acid and 0.02% heptafluorobutyric acid in a linear scheme of gradient: 0 to 6.3 minutes is 2-24% of B, then increase to 78% of B at 6.6 minutes and to 92% of B at 6.8 minutes; keep the isocratic ratio of B to 8.1 minutes following to 2% of B at 8.3 minutes. The column equilibration time at initial conditions was 3 minutes.

### Sample Preparation and Adulteration

The samples collected were stored at -20°C before the preparation. Samples were thawed at an ambient temperature and centrifuged for 5 minutes at 3,500 *g* (Eppendorf 5424R; Eppendorf AG, Germany) to sediments debris. An aliquot of 100 µL of plasma was transferred to the new clean tube and fortified with 10 µL of 10 ng/mL [^13^C_2_]-HAEE standard to 1 ng/mL concentration finally. A volume of 500 µL of methanol/acetonitrile/water at a ratio of 7:2:1 (v/v/v) was added to the sample and vortexed for 5-10 seconds. The resulting suspension was incubated at 40°C while stirring at 1,100 *g* for 10 minutes following centrifugation at 15,500 *g* and 10°C for 10 minutes to sediment proteins. Supernatant was diluted with 5 mL of cold 2.5% ammonia hydroxide in the new tube before solid-phase extraction (SPE). Solid-phase extraction was accomplished using Oasis® mixed anion-exchange (MAX) cartridges (3 cc, 60 mg of medium; Waters, Inc., Milford, MA, USA) and 20-port vacuum manifold (Waters, Inc., UK) equipped by DOA-P504A-BN pump (Gast., Inc., USA). The MAX cartridge was preconditioned consequently by 1 mL of methanol, 1 mL of acetonitrile and 1 mL of deionized water. Then the medium was washed by 2 mL of 2.5% ammonia hydroxide, and the diluted sample (5.5 mL) was loaded. After loading the sample, the medium was washed by 1mL of 2.5% ammonia hydroxide and 1 mL of deionized water following washing with 2 mL of 60% methanol in water (v/v). The endogenous HAEE and its heavy-labeled counterpart were eluted in 500 µL of 3% formic acid in acetonitrile, and the collected eluate was dried under a vacuum (Concentrator Plus; Eppendorf AG, Germany) at 30°C for about 20 minutes. The resulting dried pellet was resuspended in 35 µL of 0.1% formic acid for LS-MS analysis.

### Preparation of Standards and Calibration

Analytical [^13^C_2_]-labeled and intact (non-labeled) standards were obtained as a lyophilized powder. The stock solutions of 1 mg/mL were prepared in 30% acetonitrile/water (v/v) solution, aliquoted by 25 µL and stored at -20°C. Working solutions of 10 ng/mL were prepared by serial dilution of the stock solution in appropriate volume of 0.1% formic acid and stored at 4°C…6°C in a fridge and might be utilized for two weeks maximum after preparation. To obtain calibration curve, non-labeled standards were prepared in 1:2:5 dilution pattern to achieve a series of concentration 5 ng/mL, 2.5 mg/mL, 1 ng/mL, 0.5 ng/mL, 0.25 ng/mL, 0.1 ng/mL, 0.05 ng/mL and 0.025 ng/mL in HAEE-free matrix (Ultra-low steroid, drug-depleted DDC Mass Spect Gold® Serum; MSG3000, Sigma) fortified with the [^13^C_2_]-labeled standard in 1 ng/mL concentration. The prepared standards were analyzed in seven technical replicated, and the resulting calibration curve was tracked as a relative response in a linear regression fashion with no weighting factor applied.

### Determination of Limits

Limit of blank (LOB) has been determined using the [^13^C_2_]-labeled standards because of the endogenous nature of intact HAEE hampers to use blood plasma sample as a true negative control. To determine the limits, the obtained signal should match several identification criteria, including, signal-to-noise ratio (SNR) exceeding 10 (calculated as a root-mean-square) and retention time stability within ±1%. The accepted tolerance of relative intensities for the ion-quantifier should be within 90%-110% whereas for ion-qualifiers within ±10% (relative) for the rank-2 and rank-3 qualifiers, and ±20% by absolute for the rank-4 qualifier because its relative intensity is between 25%-50% of the ion-quantifier. Any signal of the [^13^C_2_]-labeled standards that met the identification criteria has been considered as a potential signal cut-off and threshold. Limits of detection (LOD) and quantification (LOQ) as well as the lowest limit of detection (LLOD) and limit of method (LM) were estimated based on the plotted calibration curve and recognized LOB according to the Protocols for Determination of Limits of detection and Limit of Quantitation Approved Guideline (CLSI document EP17. Wayne, PA USA: CLSI; 2004)

### Matrix Influence Factor and Recovery

The recovery was examined using plasma samples spiked with two different concentrations (1 ng/mL and 5 ng/mL) of [^13^C_2_]-labeled standards after SPE towards the corresponding standards. Matrix influence factor was calculated for the concentration of 1 ng/mL and 5 ng/mL as a ratio of difference between the signal of [^13^C_2_]-labeled standard spiked in five different plasma samples and the corresponding blank plasma samples toward the response of [^13^C_2_]-labeled standard of the corresponding concentration (1 ng/mL and 5 ng/mL) in solvent (0.1% formic acid).

### Carryover Effect

Carryover effect was inspected for three different concentrations (0.125 ng/mL, 0.5 ng/mL and 1 ng/mL) in three different schemes. The first direct scheme is supposed to run a consequent batch from least concentrated sample to the most concentrated in five technical replicates; the second scheme runs a batch in the reverse order; and the third scheme is supposed to run a batch of randomly selected samples. Before starting the observation, the reference standards (0.125 ng/mL, 0.5 ng/mL and 1 ng/mL) were analyzed in five technical replicates to reveal stability and robustness of the signal. The direct and reverse schemes covered three technical replicates for every reference concentration while the random scheme supposed technical replicates randomly organized. The carryover effect was calculated as an additive relative response from one reference standard to another.

### Inter- and Intraday Stability Assay

To examine inter- and intraday signal stability, four different plasma samples were pooled and split into nine aliquots of 100 µL. Intraday stability was assessed by preparation of three aliquots and analyzing them in three technical replicates. Inter-day stability was assessed by comparison of results obtained during three different days to highlight reproducibility of measurements between different experiments. Totally, 27 measurements were performed assuming nine measurements (three aliquots in three replicates) for three days.

### Freez-thaw Stability Assay

Several aliquots of the [^13^C_2_]-labeled standard at a concentration of 1 ng/mL in solvent (0.1% formic acid) and in human plasma were prepared to store in freezer at low temperature (-20°C) and in fridge at +4°C…+6°C. Signals were measured and monitored for 60 days to observe degradation of the substance in long-term storage conditions. To observe freeze-thawed stability, aliquots of matrix spiked with [^13^C_2_]-labeled standards at 1 ng/mL and solvent with the corresponding concentration of the standard treated by five consequent cycles of freezing and thawing during five days. Signals were measured in three replicates per cycle and the influence of interval storage was estimated by comparison of signal with the initial concentration signal before freezing. According to the recommendation of the Global Bioanalysis Consortium Harmonization Team, inspection of the signal was rescinded when the total loss of the substance reached 20% and more during the storage.

### Cross-platform Validation

Distinct aliquots of plasma samples of one hundred randomly selected samples were prepared using the proposed SPE method and the concentration of endogenous HAEE measured on QQQ mass spectrometer. Measured concentrations in each sample were compared and plotted against the results of the corresponding sample obtained previously on Q-TOF mass spectrometer. The efficiency of cross-platform transparency was fitted by the convergence between QQQ and Q-TOF measurements in terms of Pearson’s coefficient.

### Statistical Analysis and Data Handling

Raw data were processed and handled using the Mass Lynx software (Waters, Inc., version4.2 revision SCN996) if data were acquired by Q-TOF mass spectrometer, and by the Mass Hunter software (Agilent, Inc., version B10.1.67) if data were recorded on QQQ mass spectrometer. Data of the endogenous HAEE measurements among patients with Alzheimer’s disease (AD) and healthy volunteers were treated by several statistical test. The normality and homogeneity of data distribution were recognized by Kolmogorov’s and Leven’s test, respectively. Difference between the control group and different stages and AD severity was tested by ANOVA test. Validation between the healthy subjects and population of AD patients was controlled by Kruskal-Wallis and Mann-Witney non-parametric tests for independent measurements. True positive and negative samples as well as false positive and negative samples were defined using the corresponding clinical data available with plasma samples and empirical measurements. Receiver operating characteristic (ROC) and area under the curve (AUC) for the general study population were cast to determine specificity and sensitivity of the analysis and to establish recommended cut-off concentration for the positive response. Juden-index (*J*-index), positive and negative prognostic values and nomogram for determination of the positive and negative likehood ratio were calculated in Med Calc software (Med Calc Software Ltd., Belgium, version 23.0) for biological and medical statistics.

