## Supplementary figures for "Beyond Alzheimer’s story: how an engineered molecule gained an endogenous essence"

Figs. S1 to S6

**^1^H NMR and LC-UV-MS analysis data for Ac-HAEE-NH_2_ and Ac(^13^C_2_)-HAEE-NH_2_.**

Research samples of the peptides studied were synthesized by Synthon-Lab Ltd. using the Fmoc SPPS protocol on Rink-amide resin and are available as reference standards from 1MOL Ltd. (Sofia, Bulgaria). Ac-HAEE-NH_2_, Cat. no. SL104171; Ac(^13^C2)-HAEE-NH_2_, Cat. no. SL104559.

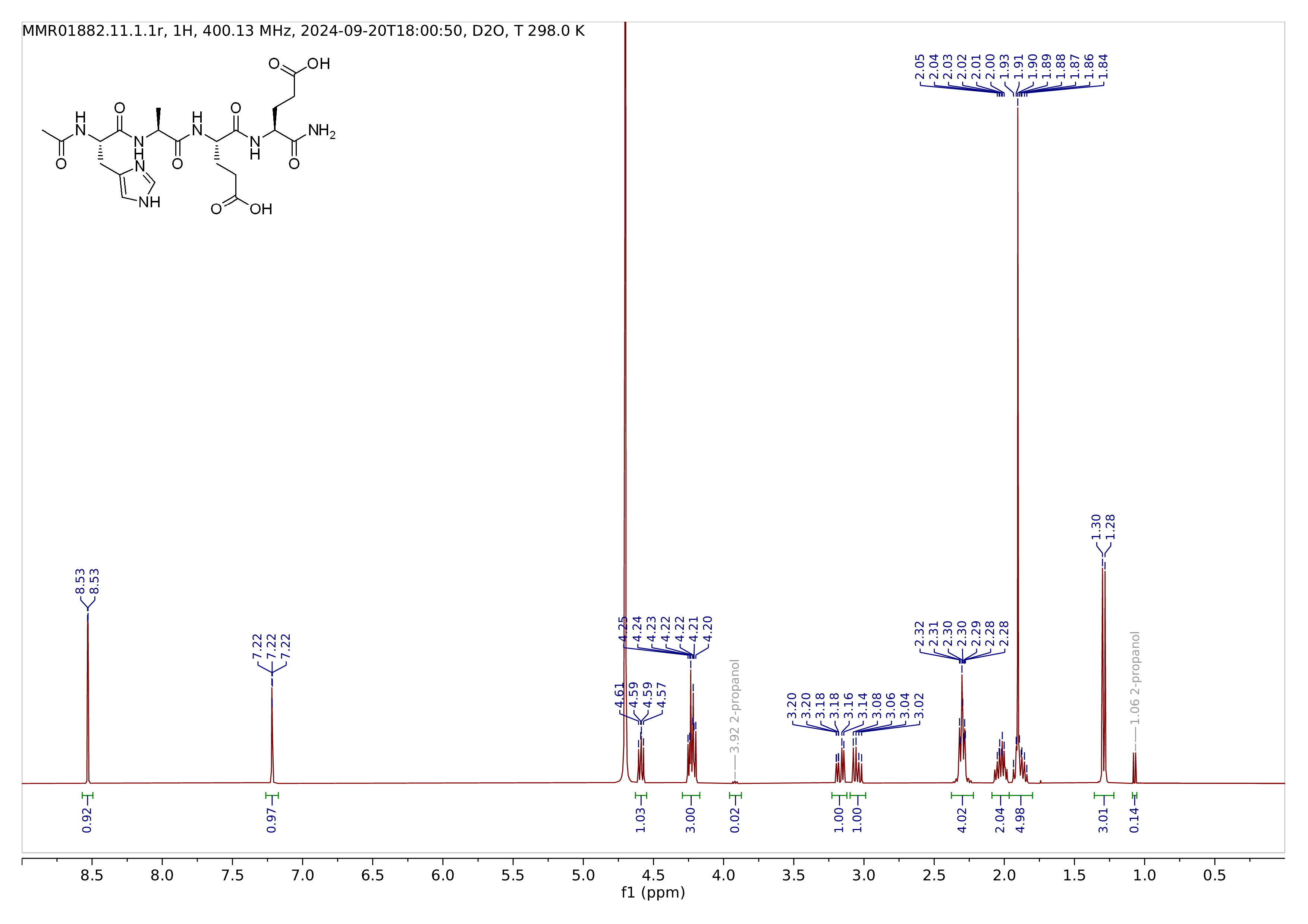

Fig. S1.

^1^H NMR of Ac-HAEE-NH_2_ (zwitterion)

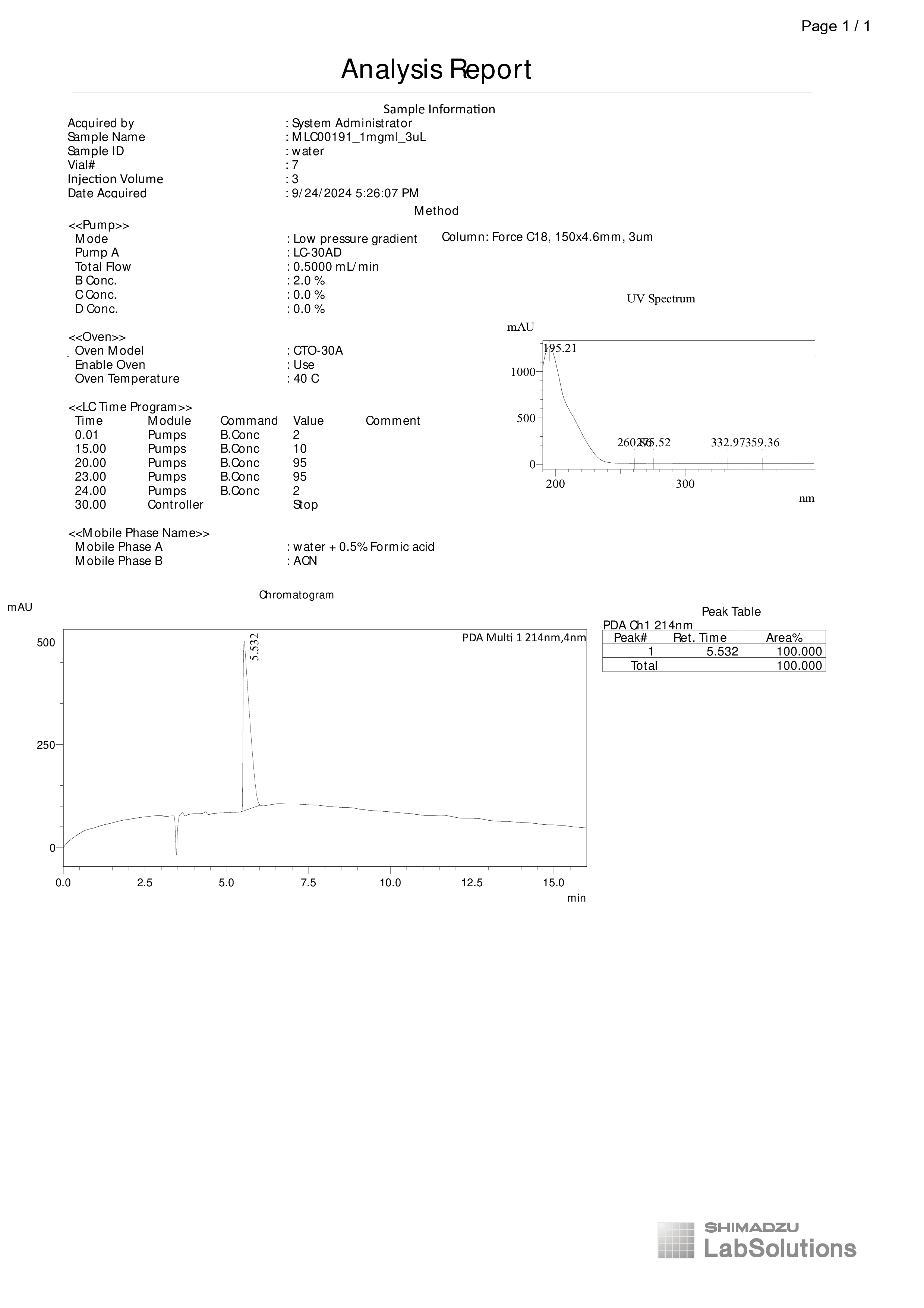

Fig. S2.

HPLC-UV of Ac-HAEE-NH_2_ (zwitterion)

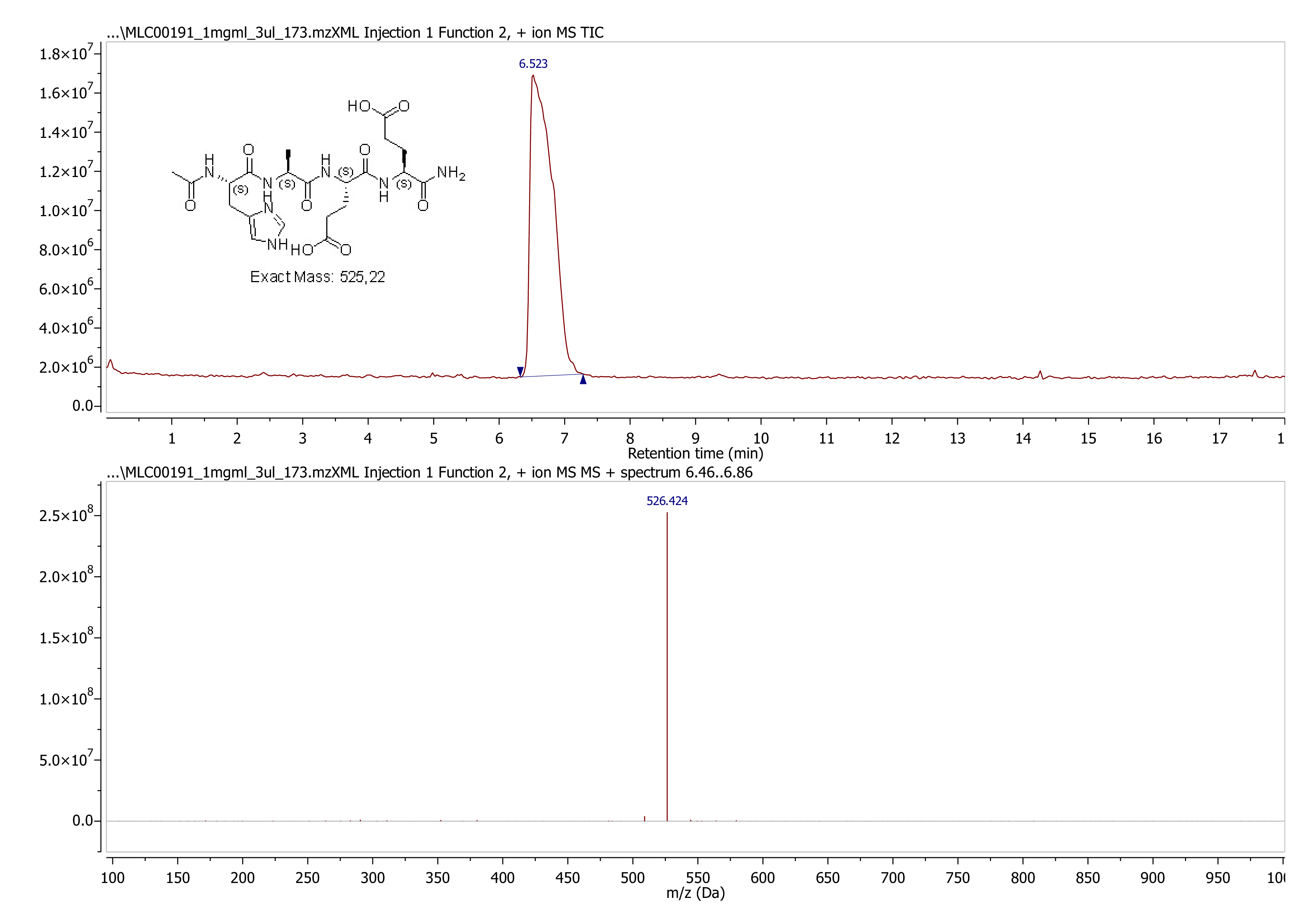
Fig. S3.

LC-MS of Ac-HAEE-NH_2_ (zwitterion)

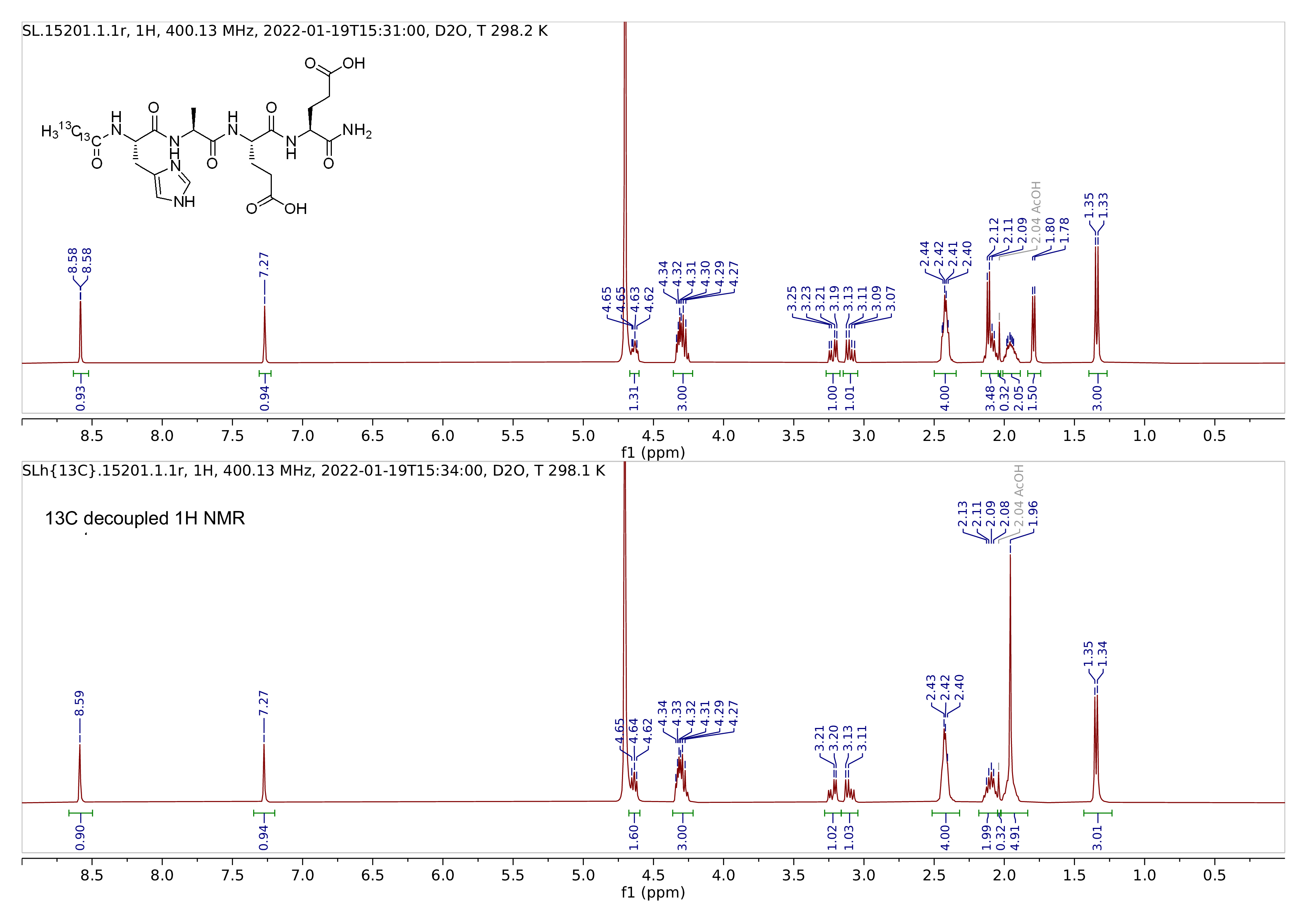
Fig. S4.

^1^H NMR of Ac(^13^C2)-HAEE-NH_2_ (TFA salt)

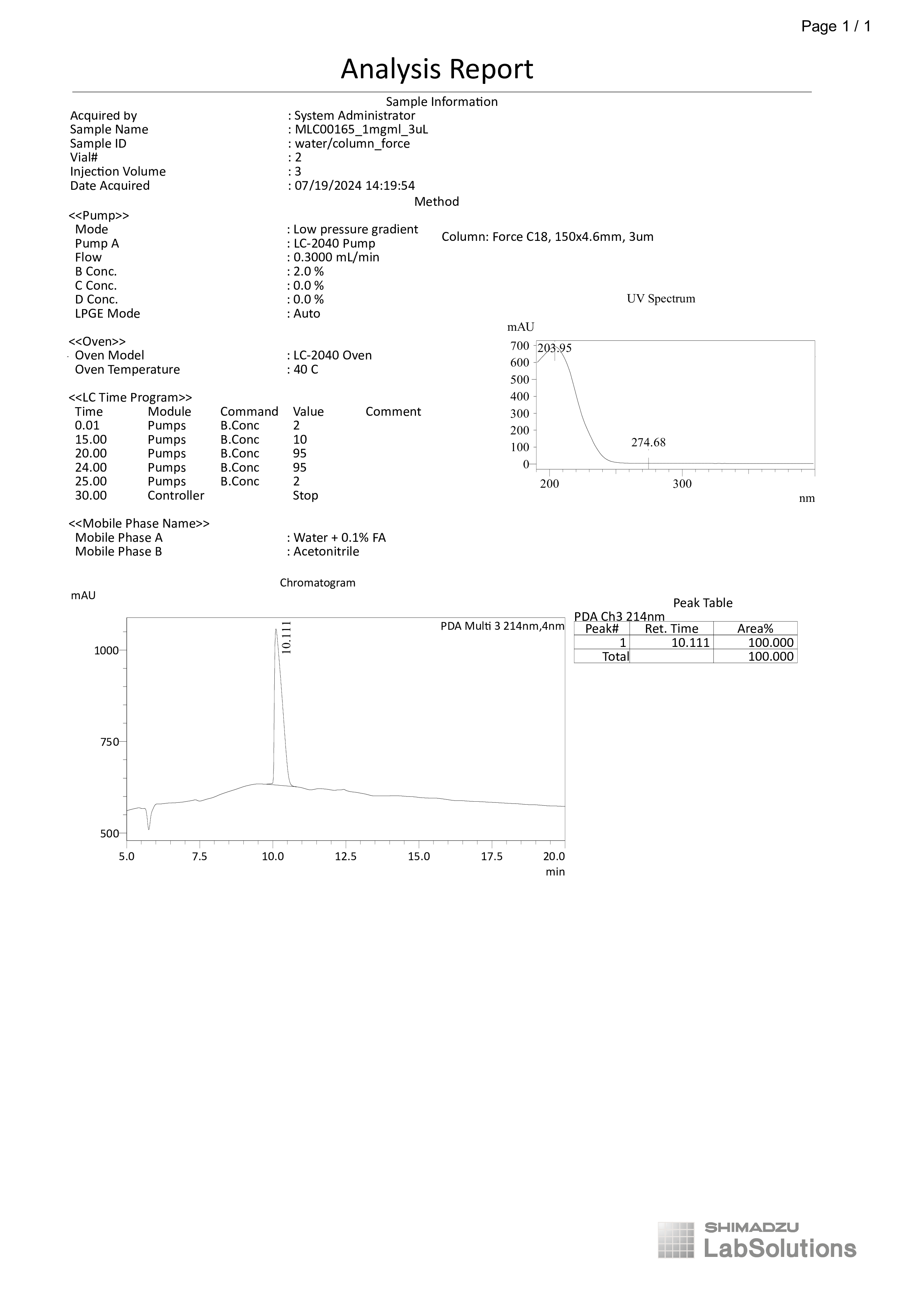
Fig. S5.

HPLC-UV of Ac(^13^C2)-HAEE-NH_2_ (TFA salt)

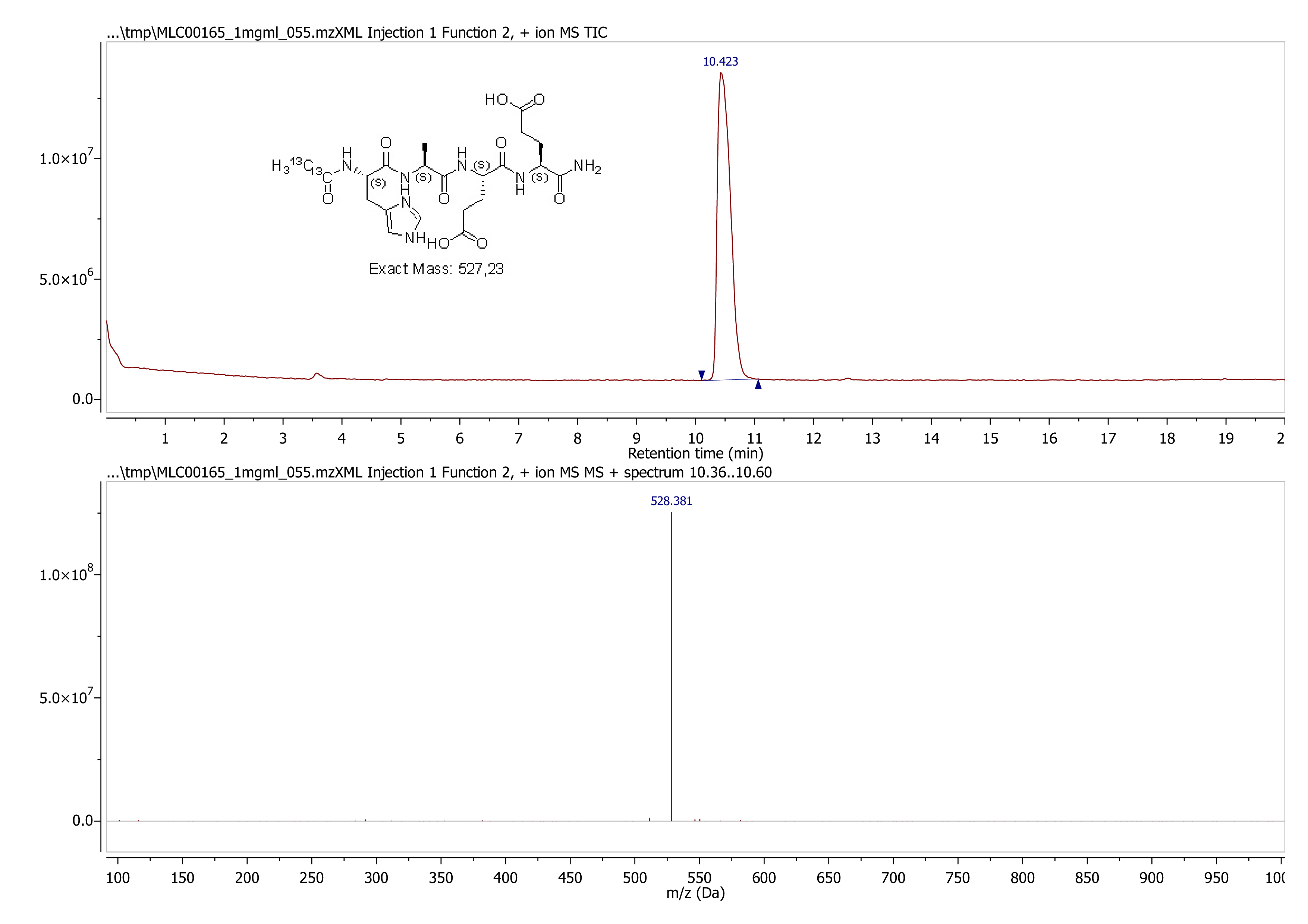

Fig. S6.

LC-MS of Ac(^13^C2)-HAEE-NH_2_ (TFA salt)
